# Isoform-specific regulation of HCN4 channels by a family of endoplasmic reticulum proteins

**DOI:** 10.1101/2020.04.10.022483

**Authors:** Colin H. Peters, Mallory E. Myers, Julie Juchno, Charlie Haimbaugh, Hicham Bichraoui, Yanmei Du, John R. Bankston, Lori A Walker, Catherine Proenza

## Abstract

Ion channels in excitable cells function in macromolecular complexes in which auxiliary proteins modulate the biophysical properties of the pore-forming subunits. Hyperpolarization-activated, cyclic nucleotide-sensitive HCN4 channels are critical determinants of membrane excitability in cells throughout the body, including thalamocortical neurons and cardiac pacemaker cells. We previously showed that the properties of HCN4 channels differ dramatically in different cell types, possibly due to the endogenous expression of auxiliary proteins. Here, we report the discovery of a family of endoplasmic reticulum transmembrane proteins that interact with and modulate HCN4. Lymphoid-restricted membrane protein (LRMP, Jaw1) and inositol trisphosphate receptor-associated guanylate kinase substrate (IRAG, Mrvi1, Jaw1L) are homologous proteins with small ER luminal domains and large cytoplasmic domains. Despite their homology, LRMP and IRAG have distinct effects on HCN4. LRMP is a loss-of-function modulator that inhibits the canonical depolarizing shift in the voltage-dependence of HCN4 activation in response to binding of cAMP. In contrast, IRAG causes a gain of HCN4 function by depolarizing the basal voltage-dependence of activation in the absence of cAMP. The mechanisms of action of LRMP and IRAG are novel; they are independent of trafficking and cAMP binding, and they are specific to the HCN4 isoform. We also found that IRAG is highly expressed in the mouse sinoatrial node where computer modeling predicts that its presence increases HCN4 availability. Our results suggest important roles for LRMP and IRAG in regulation of cellular excitability and as tools for advancing mechanistic understanding of HCN4 channel function.

**Significance statement:** The pore-forming subunits of ion channels are regulated by auxiliary interacting proteins. Hyperpolarization-activated cyclic nucleotide-sensitive isoform 4 (HCN4) channels are critical determinants of electrical excitability in many types of cells including neurons and cardiac pacemaker cells. Here we report the discovery of two novel HCN4 regulatory proteins. Despite their homology, the two proteins — lymphoid-restricted membrane protein (LRMP) and inositol trisphosphate receptor-associated guanylate kinase substrate (IRAG) — have opposing effects on HCN4, causing loss- and gain-of-function, respectively. LRMP and IRAG are expected to play critical roles in regulation of physiological processes ranging from wakefulness to heart rate through their modulation of HCN4 channel function.

## Introduction

Hyperpolarization-activated cyclic nucleotide-sensitive isoform 4 (HCN4) channels play a key role in determining membrane potential in excitable cells throughout the body. Perhaps best known as the molecular basis of the funny current (I_f_), which is critical for cardiac pacemaking in sinoatrial node myocytes (1), HCN4 channels are also important for the burst firing of thalamocortical neurons that mediates wakefulness (2, 3).

Direct binding of cAMP to a conserved, C-terminal cyclic-nucleotide binding domain (CNBD) potentiates HCN channel opening by shifting the voltage-dependence of activation to more depolarized potentials, speeding activation, and slowing deactivation (4–6). In cardiac pacemaker cells of the sinoatrial node (SAN), an increase in I_f_ in response to cAMP contributes to an increase in action potential firing rate and consequently, heart rate (7, 8). In thalamic and central neurons, HCN channel activation by cAMP depolarizes resting membrane potential, reduces membrane resistance, and increases tonic firing (3, 9). Conversely, reduction of HCN channel currents via genetic knockout, pharmacological blockers, or inhibitory regulators leads to reduced action potential firing rate and dysrhythmia in the SAN (10–12) and altered ratios of tonic and burst firing in thalamocortical neurons, which are associated with transitions between sleep and wakefulness (3, 9, 13).

We previously discovered that the cyclic nucleotide-dependent shift in the activation of HCN4 depends on the cellular context (14). When HCN4 is expressed in HEK293 cells, it exhibits the canonical depolarizing shift in voltage-dependence in response to cAMP. However, we found that when HCN4 is expressed in CHO cells, channel activation is constitutively shifted to more depolarized membrane potentials and is no longer affected by cAMP. Moreover, the constitutive activation of HCN4 in CHO cells is specific to the HCN4 isoform; HCN2 retains a large cAMP-dependent shift in voltage-dependence (14). We hypothesized that this “CHO effect” is due to expression of an endogenous, isoform-specific modulator of HCN4 and not basal phosphorylation (15) or high cAMP levels because it persists even in excised inside-out membrane patches where cAMP is absent (14) and phosphorylation is typically transient (16).

While the pore-forming subunits of ion channels can produce currents when expressed alone in heterologous systems, ion channels in native cells typically function within macromolecular complexes that include auxiliary subunits and interacting proteins that can dramatically alter channel function. For example, sodium channel β subunits regulate trafficking and inactivation gating in neurons (17); calcium channel β subunits mediate functional associations with ryanodine receptors in skeletal muscle (18, 19); and the K_v_7.1 subunit, KCNE1 (minK) is necessary for the sluggish kinetics of the slow-delayed rectifier potassium current in cardiac myocytes (20). Endogenous channels and interacting proteins in heterologous expression systems have also been shown to alter properties of transfected proteins. For example, endogenous expression of K_v_β2.1 in CHO cells changes the voltage-dependence of K_v_1.5 (21), and endogenous expression of a K_v_7 homologue in *Xenopus laevis* oocytes confounded early interpretation of the function of the minK subunit (22, 23).

Aside from cAMP, HCN channels are regulated by a plethora of other factors (24). These include filamin A, which alters HCN1 expression in neurons (25); PIP2, which potentiates opening of all HCN channel isoforms (26, 27); SAP97, which alters trafficking of HCN2 and HCN4 (28); and Src tyrosine kinase, which associates with HCN2 and HCN4 to potentiate channel opening (29, 30). One of the best studied regulators of HCN channels is the neural-specific accessory subunit, TRIP8b (31). TRIP8b interacts with HCN1, HCN2, and HCN4 at two conserved C-terminal sites to alter channel expression and decrease cAMP sensitivity (31–36). However, TRIP8b is unlikely to account for the lack of HCN4 sensitivity in CHO cells because it is not specific for the HCN4 isoform (14, 31, 32). Indeed, we did not detect endogenous TRIP8b protein expression in CHO cells in western blots (unpublished data).

In the present paper, we used the isoform-specific regulation of HCN4 in CHO cells as a starting point for the discovery of two novel HCN4-specific regulatory proteins. Using mass-spectrometry and western blotting, we identify lymphoid-restricted membrane protein (LRMP, also known as JAW1) and inositol 1,4,5-trisphoshate receptor-associated guanylate kinase substrate (IRAG, also known as MRVI1 or JAW1L) as novel HCN4 interaction partners. We show that LRMP and IRAG are HCN4 isoform-specific regulators that have opposing effects on channel gating: LRMP reduces the cAMP-sensitivity of HCN4 activation, while IRAG shifts HCN4 activation to more depolarized potentials in the absence of cAMP. In contrast to TRIP8b which competes with cAMP and changes surface expression (31, 33, 36), neither LRMP nor IRAG prevents cAMP binding or changes channel expression, suggesting that they act via novel regulatory mechanisms. We also show that IRAG is expressed at a high level in the SAN, where our computer modeling predicts that its presence increases I_f_. LRMP and IRAG are the first described HCN4 isoform-specific regulatory proteins. They are likely to play important roles in regulating cellular excitability throughout the body. And they may form a link between the ER and plasma membranes that coordinates intracellular signals and calcium release with HCN4 channel currents.

## Results

### LRMP and IRAG are HCN4 channel interaction partners

We first sought to identify novel candidate HCN4-interacting proteins that are differentially expressed in CHO and HEK cells. Silver-stained gels of HCN4 immunoprecipitates from both cell lines showed several bands that may represent cell type-specific interaction partners of HCN4 (Fig. 1A). Both the whole-cell lysate and the band at ~70 kDa in the CHO cell IP were sequenced by nano-flow reverse phase liquid chromatography mass spectrometry to identify potential interaction partners. From the candidates, lymphoid-restricted membrane protein (LRMP) was selected for further study because it appeared as a hit in both samples and because it had previously been identified in a genome-wide association study as a locus related to resting heart rate (37). Human LRMP is a 555 residue protein containing a cytoplasmic coiled-coil domain and a C-terminal transmembrane domain that is believed to anchor it to the ER membrane (Fig. 1B) (38, 39). We subsequently identified inositol trisphosphate receptor-associated guanylate kinase substrate (IRAG) as a homologue of LRMP. Human IRAG is a 904 residue protein that, like LRMP, has a cytoplasmic coiled-coil domain and a C-terminal ER membrane anchor domain (Fig. 1B) (40). IRAG has been implicated in blood pressure regulation, with one study in mice noting that IRAG knockdown leads to a decrease in resting heart rate (41). Both LRMP and IRAG have previously been shown to interact with IP3 receptors to regulate intracellular Ca^2^+ signaling (42, 43).

**Figure 1.**
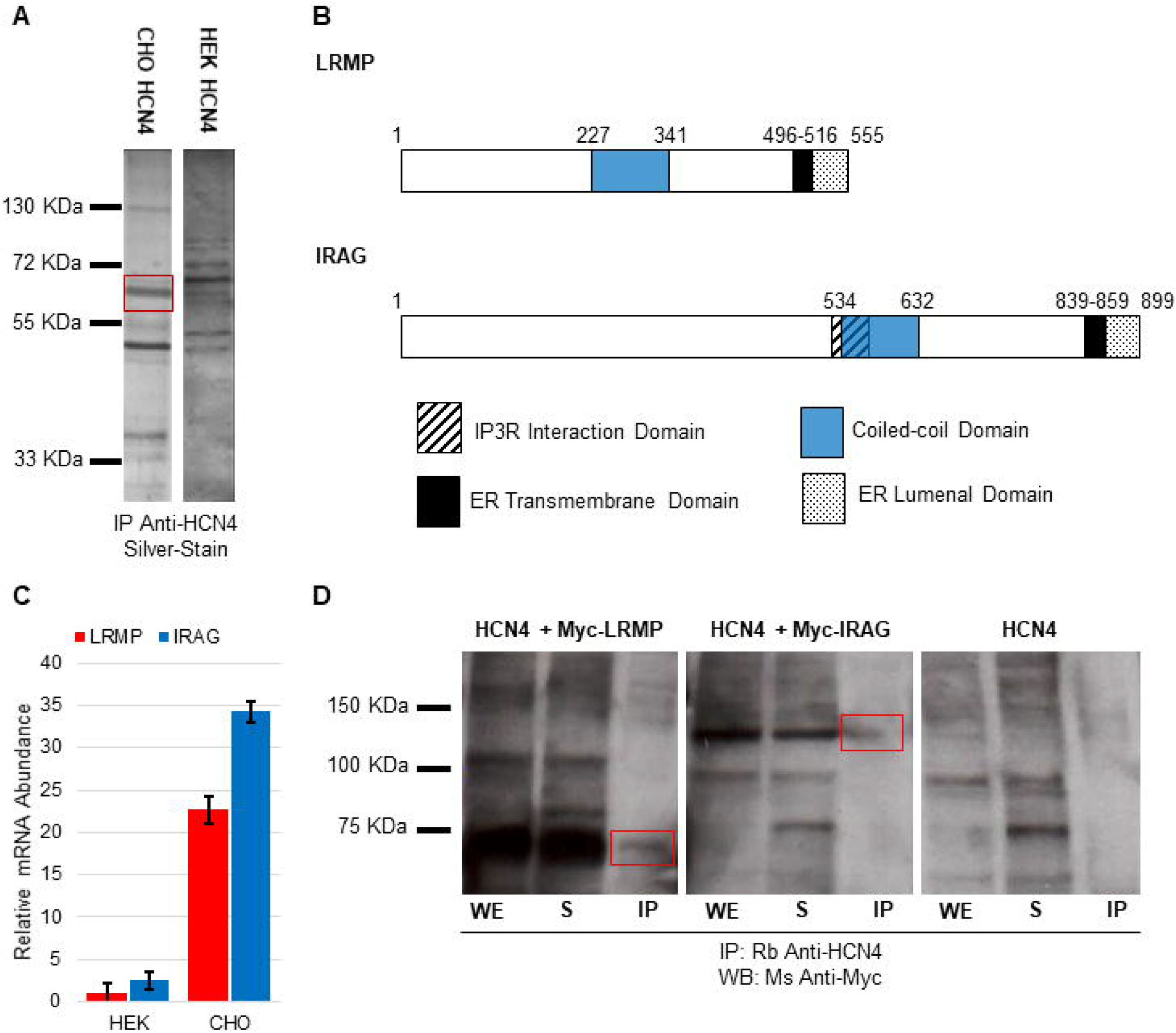
Identification of LRMP and IRAG as HCN4 interaction partners. **A**: Silver-stained gel showing proteins that were co-immunoprecipitated with HCN4 from CHO and HEK cell extracts. The box at ~70 kDa indicates a potential CHO-specific HCN4 interacting protein that was sequenced using mass-spectroscopy. **B**: Schematic illustrations of LRMP and IRAG domain structures, showing the relative sizes of the coiled-coil domains, ER transmembrane domains, ER luminal domains, and, in IRAG, the IP3 receptor interaction domain (residues 521-561). **C**: Relative mRNA abundance of LRMP *(red)* and IRAG *(blue)* in CHO and HEK cells as measured by qPCR. Data were normalized to 18S ribosomal RNA abundance and are plotted relative to LRMP abundance in HEK cells. All error bars are SEM. Data are from a minimum of 2 technical replicates of 3 independent biological samples. **D**: Western blot of anti-Myc staining of extracts of HEK 293 cells stably expressing HCN4 and transiently transfected with Myc-LRMP, Myc-IRAG, or pCDNA3.1. Red boxes show the Myc-LRMP and Myc-IRAG bands. Cell lysates were immunoprecipitated with rabbit anti-HCN4 antibodies. All panels are from the same blot with lanes removed for clarity. Representative of 3 independent blots. WE, Whole extract; S, supernatant; IP, HCN4 immunoprecipitate.

We compared the levels of endogenous LRMP and IRAG transcripts in CHO- and HEK-HCN4 stable cell lines using qPCR. Compared to HEK cells, CHO cells expressed significantly greater levels of endogenous LRMP and IRAG transcripts (P = 0.0241 and P = 0.0019, respectively; Fig. 1C; Table S1); endogenous IRAG transcript was approximately 14-fold greater in CHO cells than HEK cells and LRMP transcript was approximately 20-fold greater.

To confirm a physical association between HCN4 and both LRMP and IRAG, we transfected a HEK-HCN4 cell line with LRMP or IRAG constructs with N-terminal Myc-tags. We then immunoprecipitated HCN4 and probed the eluates with an anti-Myc antibody. Myc-LRMP and Myc-IRAG were detected in HCN4 immunoprecipitates in each of three independent experiments but no Myc labeling was seen in IPs from controls transfected with only pCDNA3.1 (Fig. 1D). We also confirmed that the anti-Myc antibody was specific for both tagged constructs and did not bind to similar weight endogenous bands (Fig. S1A). Taken together, these results show that both LRMP and IRAG associate with HCN4 and that their transcripts are upregulated in CHO cells compared to HEK cells. Based on these data and the clear differences in HCN4 function between CHO and HEK cells, we next asked whether LRMP and IRAG functionally alter currents through the HCN4 channel.

### LRMP and IRAG have opposing effects on HCN4 function

To define the functional effects of LRMP and IRAG on HCN4 channel gating, we performed whole-cell patch-clamp experiments in HEK cells stably expressing HCN4 (Table S2). Representative HCN4 currents in the absence or presence of LRMP or IRAG, and with and without 1 mM cAMP are shown in Fig. 2A. As expected, addition of 1 mM cAMP to the recording pipette in the absence of LRMP or IRAG significantly shifted the midpoint activation voltage (V_1/2_) of HCN4 channels to more depolarized membrane potentials, as assessed by Boltzmann fits of average conductance-voltage relations (P < 0.0001; Fig. 2B-D).

**Figure 2.**
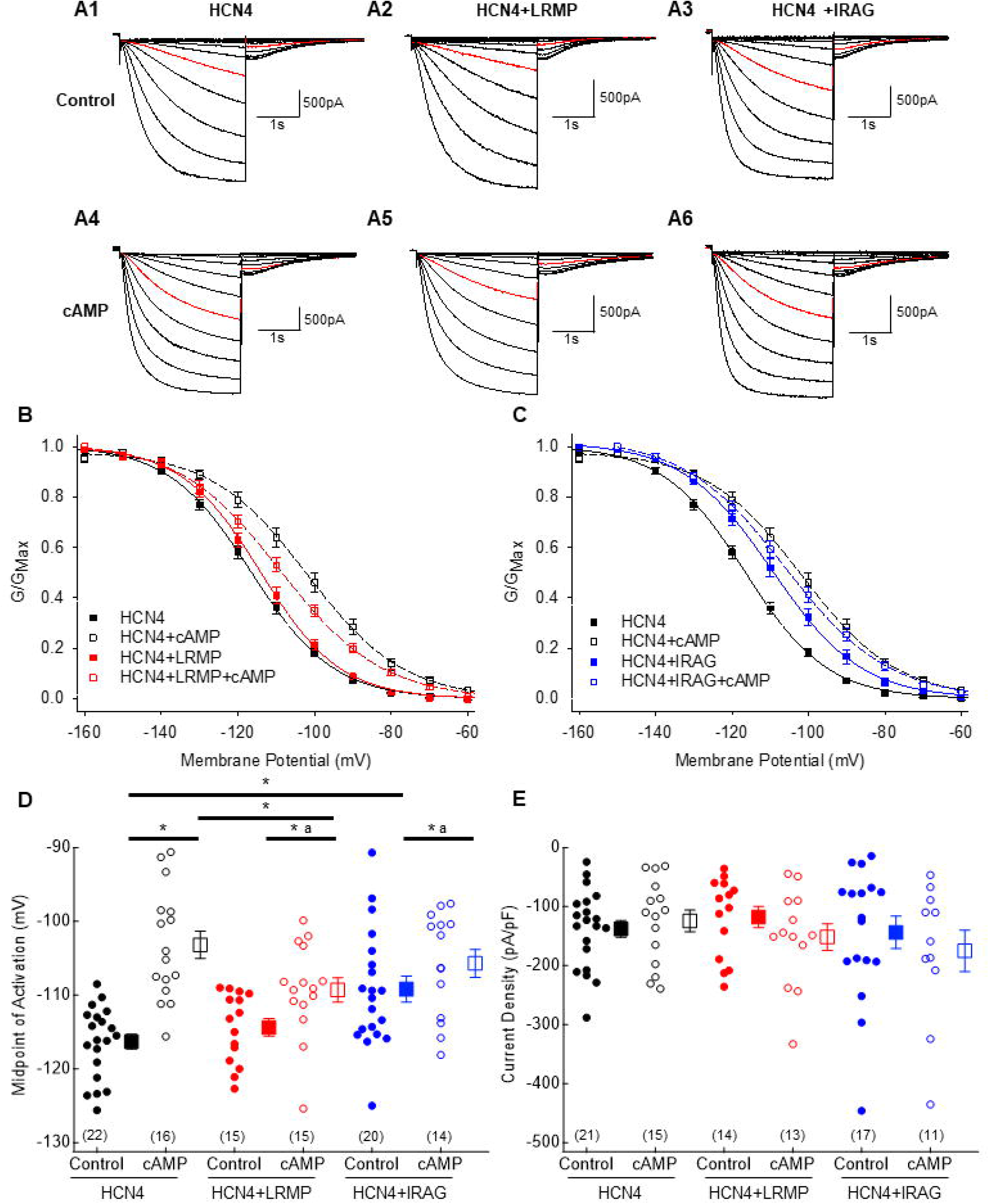
LRMP and IRAG have opposing effects on HCN4 function. **A1-6**: Representative whole cell HCN4 currents from HEK cells in the absence or presence of LRMP or IRAG with or without 1 mM cAMP in the patch pipette. Currents were elicited with 3 s hyperpolarizations to membrane potentials between −50 mV and −150 mV in 10 mV increments followed by 3 s pulses to −50 mV. Red traces are the currents at −110 mV. **B, C:** Average conductance-voltage relations for HCN4 in control conditions *(black),* the presence of LRMP *(red),* or the presence of IRAG *(blue).* GVs in the presence of 1 mM cAMP are shown by *open symbols.* Error bars in this and subsequent panels are SEM, N = 14-22 (See panel **D**). Control HCN4 data in panel **C** are the same as those in panel **B. D**: Average V_1/2_ values for HCN4 in HEK cells in the absence or presence of LRMP *(red)* or IRAG *(blue)* and 1 mM cAMP *(open).* Each individual observation is plotted as a *circle.* Number of observations for each dataset is given in parentheses. Averages (± SEM) are plotted as *squares.* **E**: Average current density in response to a 3 s step to −150 mV of HCN4 in HEK cells in the absence or presence of LRMP, IRAG, and cAMP using the same color scheme as **D**. * indicates P<0.05 between two means (see text for P-values). ^a^indicates that the cAMP-dependent shift in the presence of LRMP or IRAG is significantly different than the corresponding shift in control.

In the absence of cAMP, LRMP had no effect on the V_1/2_ of HCN4 (P = 0.3709; Fig. 2B and 2D). However, the presence of LRMP reduced the cAMP-dependent shift in channel activation; with 1 mM cAMP in the patch pipette, the V_1/2_ of HCN4 was significantly more hyperpolarized in the presence of LRMP versus the absence of LRMP (P = 0.0113; Fig. 2B and 2D). Thus, LRMP causes a loss-of-function (LOF) of HCN4 by reducing the cAMP-dependent shift in the V_1/2_ from ~13 mV to ~5 mV. In contrast to LRMP, IRAG significantly shifted the basal V_1/2_ by approximately 7 mV to more depolarized potentials in the absence of cAMP (P = 0.0006; Fig. 2C and 2D). And IRAG expression did not significantly change the V_1/2_ in the presence of cAMP (P = 0.2978; Fig. 2C and 2D). Thus, like LRMP, the presence of IRAG reduces the cAMP dependent shift in HCN4 to ~5 mV; however, in the case of IRAG the reduction is due to an IRAG-dependent gain-of-function (GOF) that pre-shifts the V_1/2_ to more depolarized potentials in the absence of cAMP. We obtained similar results with N-terminal Myc-tagged LRMP and IRAG constructs, confirming the above findings and demonstrating that a small N-terminal tag does not interfere with the ability of either LRMP or IRAG to regulate HCN4 (Fig. S2).

Importantly, neither LRMP nor IRAG significantly altered the current density of HCN4 at −150 mV in either the absence (P = 0.5021 and P = 0.8307, respectively; Fig 1E) or presence (P = 0.4057 and P = 0.1436, respectively; Fig 1E) of cAMP. Thus, while LRMP and IRAG modulate HCN4 gating, they differ from the well-studied HCN4 accessory protein, TRIP8b, which alters channel trafficking to the membrane (31, 33–35), suggesting that they act via different sites and mechanisms.

### LRMP and IRAG do not prevent cAMP binding to the CNBD

To gain further insight into the mechanisms of action of LRMP and IRAG, we examined the ability of cAMP to increase the rate of activation and decrease the rate of deactivation of HCN4 as a proxy for the ability of cAMP to bind to the CNBD. We determined the time course of HCN4 activation by measuring the time to half-maximal current during hyperpolarizing pulses to −150 mV and fit the deactivation time course at −50 mV to an exponential decay to determine deactivation time constants (Table S3). As expected, 1 mM cAMP significantly sped the rate of channel activation for HCN4 alone at −150 mV (P < 0.0001; Fig. 3A1 and 3B). This effect was preserved, albeit diminished, in cells transfected with LRMP or IRAG (P = 0.0212 and P = 0.0214, respectively; Fig. 3A2, 3A3, and 3B). Transfection of LRMP did not significantly alter the kinetics of activation at −150 mV in the absence or presence of cAMP compared to control (P = 0.1390 and P = 0.0836, respectively). Transfection of IRAG significantly accelerated channel activation at −150 mV compared to control in the absence of cAMP, but not when 1 mM cAMP was present in the recording pipette (P = 0.0048 and P = 0.6079). The increase in the rate of activation in the presence of IRAG and absence of cAMP was explained by the +7 mV shift in the basal V_1/2_ of IRAG-transfected cells (Fig. 2C). When the rates of activation measured in IRAG-transfected cells were shifted to 7 mV more hyperpolarized potentials to account for the shift in midpoint, the predicted time to half-maximal activation at −150 mV was 398 ms, similar to the 372 ms in the absence of IRAG.

**Figure 3:**
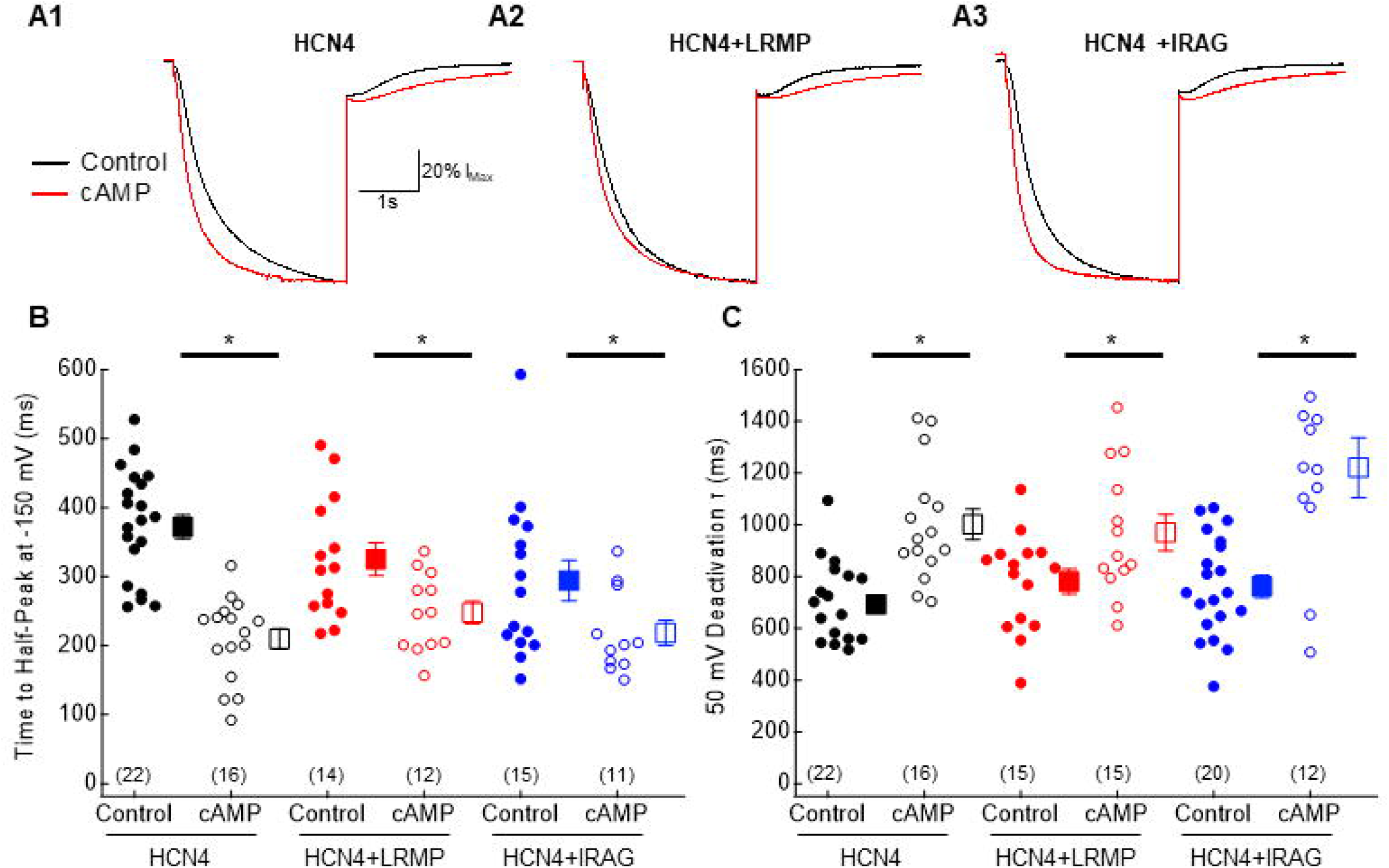
LRMP and IRAG do not prevent cAMP binding to the CNBD. **A1-3:** Representative current traces of HCN4 in the absence or presence of LRMP or IRAG and the absence *(black)* or presence of 1 mM cAMP *(red).* Currents were elicited with 3 s hyperpolarizations to −150 mV followed by 3 s pulses to −50 mV. **B:** Average time to half maximal current at −150 mV of HCN4 in control *(black)* or presence of LRMP *(red)* or IRAG *(blue)* and 1 mM cAMP *(open).* Error bars in this and **C** are SEM. Each individual observation is plotted as a *circle* and averages are plotted as *squares.* Number of observations for each dataset are given in parentheses. **C:** Average deactivation time constant of HCN4 at −50 mV in the absence or presence of LRMP, IRAG, and cAMP using the same color scheme as **B**. 4 outliers are not shown in **C**. * indicates P<0.05 between two means (see text for P-values).

We next examined cAMP-dependent slowing of deactivation. As expected, the presence of cAMP significantly slowed the deactivation time constant for HCN4 at −50 mV (P < 0.0001; Fig. 3A1 and 3C). A similar, significant cAMP-dependent slowing of deactivation was still present when either LRMP or IRAG was transfected (P = 0.0028 and P < 0.0001, respectively; Fig. 3A2, 3A3, and 3C). Neither LRMP nor IRAG affected the rate of deactivation at −50 mV in the absence of cAMP (P = 0.2619 and P = 0.3492, respectively; Fig. 3C) or the presence of cAMP (P = 0.8234 and P = 0.3051, respectively; Fig. 3C). These data indicate that LRMP and IRAG do not act by preventing cAMP binding to the CNBD.

### LRMP and IRAG are isoform-specific modulators of HCN4

We next asked whether LRMP or IRAG are specific for the HCN4 isoform or whether they can also regulate HCN1 or HCN2 channels. We performed whole-cell patch-clamp experiments on HEK cells transiently transfected with either HCN1 or HCN2 (Table S4). Neither LRMP (Fig. 4A) nor IRAG (Fig. 4C) significantly altered the V_1/2_ of channel activation of HCN1 in the absence of cAMP (P = 0.3579 and P = 0.2411, respectively; Fig. 4E). As previously reported, the presence of 1 mM cAMP in the recording pipette did not significantly shift the midpoint of activation of HCN1 (44), and the channels remained insensitive to cAMP in the presence of LRMP or IRAG (P ≥ 0.2694 for all conditions; Fig. 4E). As expected, the presence of 1 mM intracellular cAMP dramatically shifted HCN2 activation towards more depolarized potentials (P < 0.0001; Fig. 4F). The presence of LRMP (Fig. 4B) or IRAG (Fig. 4D) did not change this sensitivity or alter the V_1/2_ of HCN2 in the absence or presence of cAMP (P > 0.3715 for all conditions; Fig. 4F). These data indicate that both LRMP and IRAG are specific for the HCN4 isoform, reminiscent of the “CHO effect” where HCN4 channels are insensitive to cAMP, while HCN2 channels respond normally.

**Figure 4.**
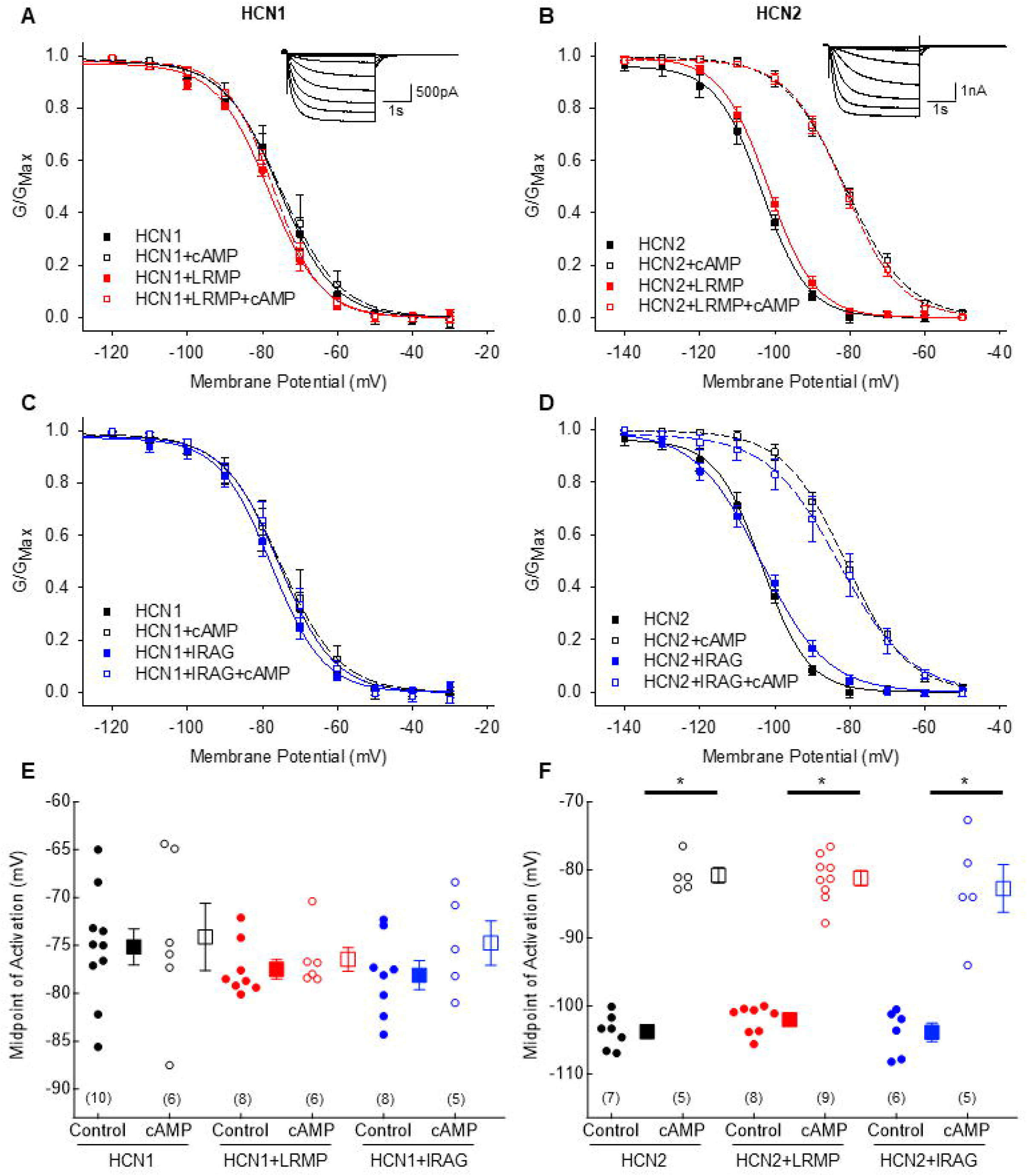
LRMP and IRAG are isoform-specific modulators of HCN4. **A, C:** Average conductance-voltage relationships for HCN1 in control conditions *(black),* the presence of LRMP *(red),* or the presence of IRAG *(blue).* GVs in the presence of 1 mM cAMP are shown by *open symbols.* Error bars in this and subsequent panels are SEM, N = 5-10 (See panel **E**). Control HCN1 data in panel **C** are the same as those from panel **A**. *Inset:* Representative currents of HCN1 elicited with 3 s hyperpolarizations to membrane potentials between −30 mV and −130 mV followed by a 3 s pulse to −50 mV. **B, D:** Average conductance-voltage relationships for HCN2 in the absence or presence of LRMP, IRAG, and cAMP using the same color scheme as **A**. N = 5-9 (See panel **F**). Control HCN2 data in panel **D** are the same as those from panel **B**. *Inset:* Representative currents of HCN2 elicited with 3 s hyperpolarizations to membrane potentials between −50 mV and −150 mV followed by a 3 s pulse to −50 mV. **E**: Average V_1/2_ values for HCN1 in HEK cells in the absence or presence of LRMP *(red)* or IRAG *(blue)* and 1 mM cAMP *(open).* Each individual observation is plotted as a *circle* and averages are plotted as *squares.* Number of observations for each dataset are given in parentheses. **F**: Average V_1/2_ values for HCN2 in HEK cells in the absence or presence of LRMP, IRAG, and cAMP using the same color scheme as **E**. * indicates P<0.05 between two means (see text for P-values).

### Endogenous LRMP is responsible for the lack of cAMP sensitivity in CHO cells

As LRMP was detected as a potential endogenous HCN4 interacting proteins in CHO cells, we used CRISPR knock-down of LRMP and IRAG to determine whether they are sufficient to account for the “CHO effect” (i.e. the depolarized basal V_1/2_ and lack of cAMP sensitivity of HCN4 in CHO cells (14); Table S4). Transfection of CHO cells with a control CRISPR without a gRNA did not alter HCN4 channel activation in either the absence or presence of cAMP, and the channels remained insensitive to cAMP in the CHO cell context (P = 0. 1886 and P = 0.2412, respectively; Fig. 5A). CRISPR-mediated knock-down of endogenous LRMP in CHO cells caused a significant hyperpolarizing shift in the midpoint of HCN4 activation in the absence of cAMP compared to cells transfected with the blank CRISPR plasmid (P = 0.0046; Fig. 5B and 5D). Furthermore, the knockdown of LRMP in CHO cells restored a significant, ~13.2 mV depolarizing shift in the V_1/2_ for HCN4 in response to cAMP (P < 0.0001; Fig. 5B and 5D). The magnitude of shift with LRMP knock-down is similar to the shift seen in HEK cells (Fig. 2A). In contrast, transfection of plasmids to knock-down endogenous IRAG did not significantly shift the V_1/2_ for HCN4 compared to cells transfected with the blank CRISPR plasmid in either the absence or presence of cAMP (P = 0.4605 and P = 0.7603, respectively; Fig. 5C and 5D). Thus, endogenous LRMP in CHO cells not only reduces the cAMP-dependent shift in HCN4 activation, as is seen in HEK cells, but also shifts the V_1/2_ to more depolarized potentials in the absence of cAMP. These data suggest that LRMP accounts for the majority of the “CHO effect” on HCN4.

**Figure 5.**
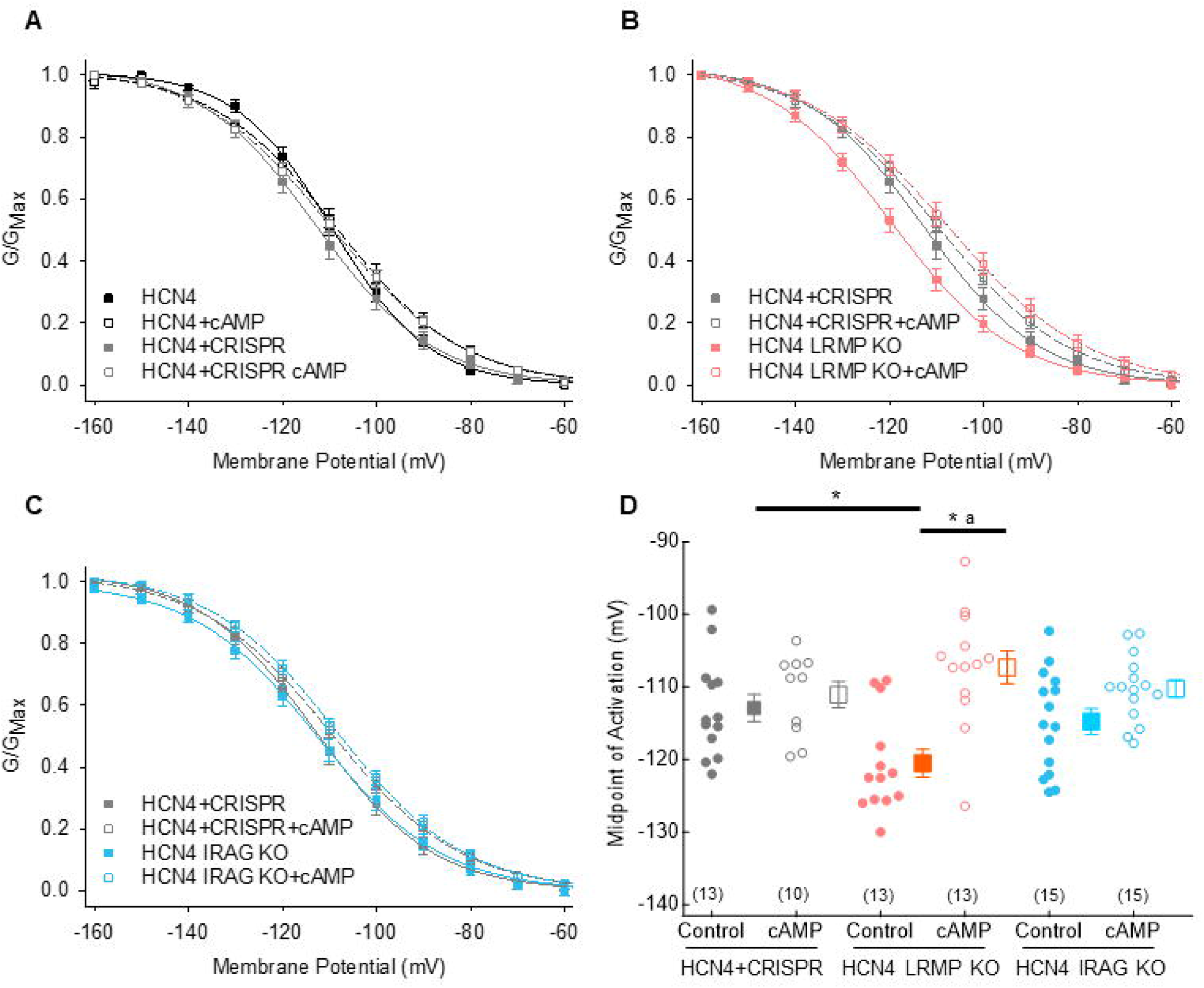
Endogenous LRMP is responsible for the lack of cAMP sensitivity in CHO cells. **A-C:** Average conductance-voltage relationships for HCN4 in CHO cells in control conditions *(black),* the presence of CRIPSR Cas9 *(grey),* the presence of CRISPR Cas9 and gRNAs targeted to CHO LRMP *(red),* or the presence of CRISPR Cas9 and gRNAs targeted to CHO IRAG *(blue).* GVs in the presence of 1 mM cAMP are shown by *open symbols.* Error bars in all panels are SEM, N = 10-15 (See panel **D**). HCN4 CRISPR control data for panels A-C are the same. **D:** Average V_1/2_ values for HCN4 in CHO cells in the presence of CRISPR Cas9 and the absence or presence of gRNAs targeted to LRMP *(red)* or IRAG *(blue)* and 1 mM cAMP *(open).* Each individual observation is plotted as a *circle* and averages are plotted as *squares.* Number of observations for each dataset are given in parentheses. * indicates P<0.05 between two means (see text for P-values). ^a^indicates that the cAMP-dependent shift when gRNAs targeted to LRMP are present is significantly different than the corresponding shift when only CRISPR Cas9 is present.

### LRMP & IRAG transcript and IRAG protein are expressed in mouse sinoatrial node tissue

As an indication of the potential role for LRMP or IRAG modulation of HCN4 in native cells, we next assessed whether LRMP or IRAG are expressed in the SAN, which is known to have high expression of HCN4. We used qPCR to measure transcript levels of HCN4, LRMP, and IRAG in SAN tissue (Fig. 6A). We found that IRAG is abundantly expressed in the SAN, at levels not significantly different from those of HCN4 (P = 0.4569). LRMP transcript was expressed at non-negligible levels, but the mRNA abundance was significantly lower than for HCN4 or IRAG (P < 0.0001 and P = 0.0005, respectively).

**Figure 6:**
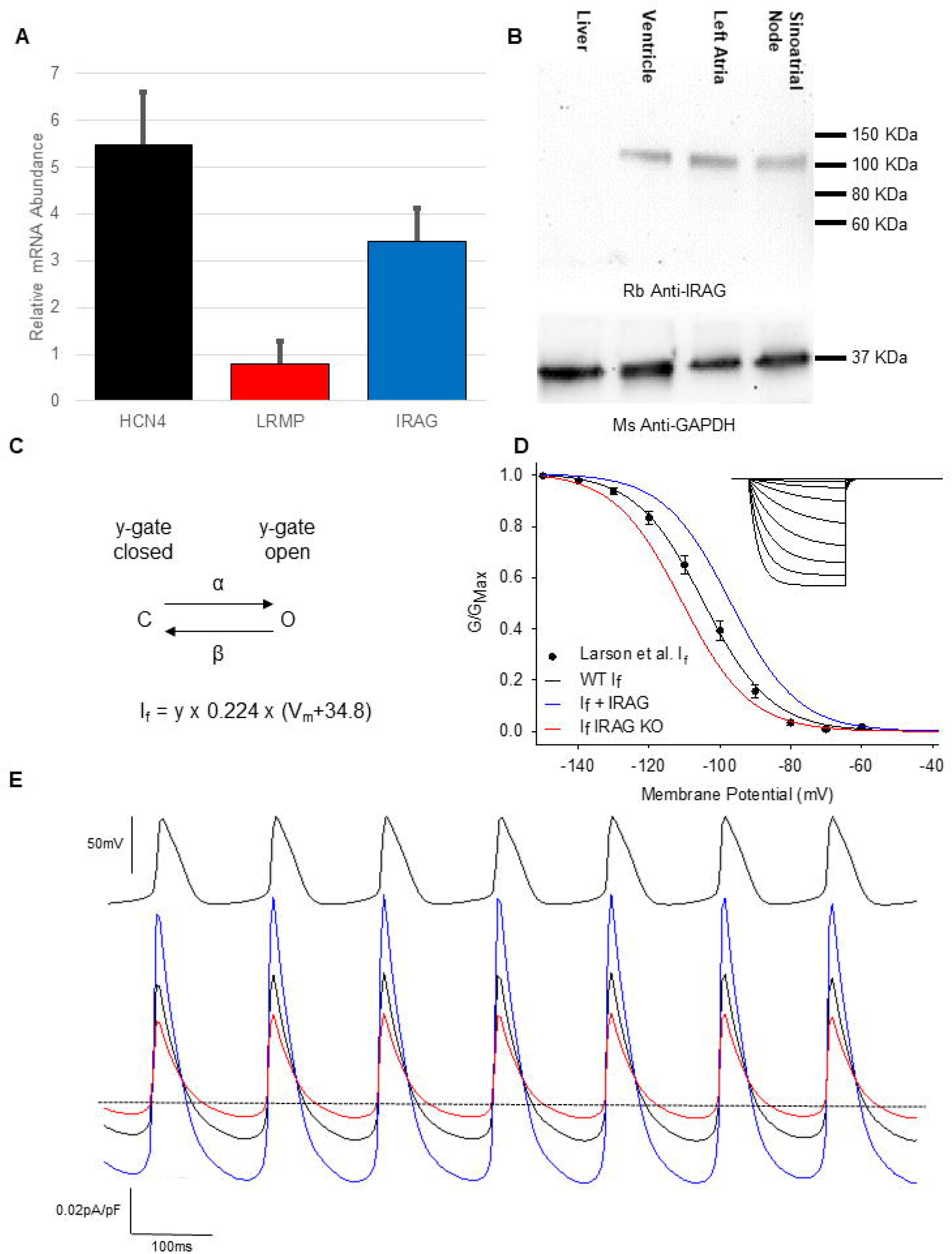
LRMP & IRAG are expressed in mouse sinoatrial node tissue and IRAG is predicted to increase I_f_ during sinoatrial APs. **A:** Relative mRNA abundance of HCN4 *(black),* LRMP *(red),* and IRAG *(blue)* in mouse sinoatrial node tissue as measured by qPCR. In all cases, data were normalized to 18S ribosomal RNA abundance and are plotted relative to HCN4 abundance in left atrial tissue from the same mice. Error bars are SEM. Data are from a minimum of 4 technical replicates of 3 independent biological samples. **B:** Western blot of IRAG in lysates from mouse sinoatrial node, left atrium, ventricle, and liver. GAPDH is shown as a loading control. The blot is representative of 3 independent biological samples. **C:** Schematic of the I_f_ model developed by Verkerk and Wilders (46). See methods for equations used in the model. **D:** Simulated voltage-dependence of I_f_ in the wild-type *(black lines),* IRAG overexpression *(blue lines),* and IRAG knockout models *(red lines)* overlaid on experimental data from Larson *et al. (black symbols)* collected in young mice with basal levels of β-adrenergic stimulation (47). *Inset:* Simulated wild-type I_f_ currents during 3 s hyperpolarizing pulses to membrane potentials between −50 mV and −150 mV followed by a 3 s pulse to −50 mV. **E:** Simulated I_f_ currents in the wild-type *(black)*, IRAG overexpression *(blue),* and IRAG knockout models *(red)* stimulated with a train of action potentials recorded from a mouse sinoatrial node myocyte (top).

We then assessed IRAG protein expression in isolated SAN tissue by western blotting. Liver protein was used as a negative control based on the low level of mRNA reported in previous RNASeq datasets in the Human Protein Atlas (45). We validated the anti-IRAG antibody by showing that it specifically bound to mouse IRAG and not to endogenous proteins in transfected HEK cells (Fig. S1B). Unfortunately, the commercially available LRMP antibodies were found to be non-specific. IRAG protein expression was found to be similar in homogenates of mouse SAN tissue, left atrial tissue, and ventricular tissue, but was absent in the liver (Fig. 5B). IRAG, therefore, may be capable of modulating HCN4 function in the SAN where both proteins are abundantly expressed.

### IRAG expression is predicted to increase I_f_ in sinoatrial myocytes

As IRAG leads to an increase in HCN4 activity and is expressed within the SAN, we used a previously proposed model of I_f_ (Fig. 6C) to predict the effect of IRAG expression on I_f_ in mouse sinoatrial myocytes (46). We scaled the time constants and steady-state activation of the model to fit conductance-voltage relationships of I_f_ from mouse sinoatrial myocytes previously reported by our lab (Fig. 6D) (47). We then created two models to predict the range of effects that IRAG could have on I_f_. IRAG over-expression or IRAG knockout were incorporated as 6-7 mV voltage shifts in the steady-state activation and time constants of activation and deactivation to replicate the magnitude of effect seen in HEK cells (see methods). When stimulated with action potential waveforms previously recorded from mouse sinoatrial myocytes in our lab, the simulated IRAG over-expression increased both the inward I_f_ active during the diastolic depolarization phase and the outward I_f_ during the AP upstroke, while the simulated knock-out of IRAG reduced I_f_ (Fig. 6E). The physiological effects of IRAG are likely to be bounded by this range. As we have already shown that IRAG is expressed in the sinoatrial node (Fig. 6A and 6B), we expect that it contributes to a relatively depolarized V_1/2_ of I_f_ in sinoatrial myocytes, consistent with studies in an IRAG knockdown mouse that show a decrease in resting heart rate (41). Since I_f_ is an important determinant of the diastolic depolarization rate and heart rate (48), these data predict that IRAG plays a physiologically relevant role in defining the magnitude of I_f_ and, consequently, heart rate.

## Discussion

In this paper we identify LRMP and IRAG as two novel, isoform-specific modulators of HCN4. Although LRMP and IRAG are homologues and both physically associate with HCN4, they exert opposing effects on the channels. LRMP causes a loss of HCN4 function by reducing the cAMP-dependent shift in the voltage-dependence of activation whereas IRAG causes a GOF by shifting the activation of HCN4 to more depolarized potentials in the absence of cAMP. Moreover, LRMP and IRAG appear to act by unique mechanisms compared to other HCN channel regulators such as TRIP8b, in that they do not compete with cAMP for binding to the CNBD, affect current density, or regulate other HCN isoforms. We also found that IRAG is expressed at high levels in the SAN where its regulation of HCN4 is predicted to play a physiologically relevant role in modulating I_f_.

### LRMP is the CHO factor

Consistent with our original identification of LRMP in CHO cell immunoprecipitates (Fig. 1), we found that LRMP transcript is expressed at much higher levels in CHO cells than in HEK cells. When LRMP is knocked out using CRISPR Cas9 and targeted gRNAs, the “CHO effect” is removed and HCN4 again responds normally to cAMP. This indicates that LRMP is a key factor in disrupting the cAMP-dependent shift in HCN4 activation in CHO cells. The lack of effect from the IRAG knockout in CHO cells was surprising given that it is expressed as abundantly as LRMP at the transcript level; it is possible that the IRAG transcript expression in CHO cells does not correlate with protein expression (49). In total, these data underscore the point that heterologous expression systems are not blank slates. Endogenous expression of subunits and regulators can cause dramatic differences in ion channel function between expression systems (20, 21), in much the same way that they do between cell types within the body.

Importantly, our identification of LRMP does not preempt the existence of other endogenous regulators of HCN4 in CHO cells. Our immunoprecipitation experiments (Fig 1A) also show bands at ~35, 55, and 130 kDa we have yet to account for. Furthermore, our results do not explain why LRMP causes a shift in the basal V_1/2_ of HCN4 in CHO cells in addition to the decrease in sensitivity to cAMP that is also seen in HEK cells. It is possible that this effect is due to the presence of other endogenous proteins or differences in the phosphorylation status of HCN4 or associated proteins in CHO and HEK cells. Indeed, preliminary results from our lab suggest some differences in the response of HCN4 to alkaline phosphatase between CHO and HEK cells (50).

The stoichiometry between LRMP and HCN4 will also need to be assessed in the future. As the endogenous LRMP in CHO cells is likely present at a lower level than the over-expressed HCN4, we hypothesize that LRMP may be able to exert its effects with less than 4 LRMP subunits per channel. Unfortunately, the lack of a specific LRMP antibody prevents us from evaluating the endogenous protein level or the degree of knockdown at this time.

### LRMP & IRAG have opposing effects

Despite the fact that LRMP and IRAG both immunoprecipitate with and modulate HCN4, they do not act in the same manner. LRMP causes a LOF by reducing the depolarizing shift in V_1/2_ caused by cAMP while IRAG causes a GOF by shifting channel activation to more positive potentials in the absence of cAMP. One possible model is that both homologues bind HCN4 via a common interaction site that tethers LRMP and IRAG to the channel. In such a model, LRMP and IRAG would then exert opposing effects on channel gating via different effector sites. While LRMP and IRAG share coiled-coil and ER-transmembrane domains, they diverge considerably in the length and sequence of the cytoplasmic domains that are N-terminal to the coiled-coil domain (Fig. 1B). Interestingly, although small Myc tags were tolerated on the N-terminals of LRMP and IRAG (Fig. S2), when a larger GFP tag was added to the N-terminus of LRMP, the protein expressed, but no longer had a functional effect on HCN4 (data not shown). Based on these observations, we hypothesize that LRMP and IRAG exert their effects on HCN4 via different sites that lie towards the N-terminal domains of the proteins.

### LRMP and IRAG likely interact with unique sequences in distal N- and/or C-terminals of HCN4

Although LRMP and IRAG alter the cAMP-dependent shifts in the voltage-dependence of HCN4 activation, cAMP still slows the channel deactivation rate (Fig. 3). This indicates that cAMP can still bind to the CNBD even in the presence of LRMP or IRAG. The lack of competition with cAMP contrasts LRMP and IRAG to the well-known HCN channel accessory protein, TRIP8b, which decreases cAMP sensitivity in all HCN isoforms through a mechanism that is at least partially competitive and involves direct binding of TRIP8b to the CNBD (32, 34–36). In the case of TRIP8b, these effects are further stabilized by a second interaction at a conserved SNL sequence in the C-terminus that, depending on splice variant, can increase or decrease HCN channel expression on the plasma membrane (33–35). This marks another difference between TRIP8b and LRMP and IRAG, which do not alter channel expression (Fig. 2E).

Importantly, LRMP and IRAG also differ from TRIP8b in that they are specific for the HCN4 isoform (31–33). Consistent with our previous data showing unaltered HCN2 function in CHO cells (14), neither LRMP nor IRAG alters the function of HCN1 or HCN2 (Fig. 4). In addition to the lack of competition with cAMP, the isoform-specificity of LRMP and IRAG suggests that the binding site on HCN4 for these proteins is not likely to be within the CNBD, the transmembrane region, or the HCN domain in the proximal N-terminus, because these regions are all highly conserved across isoforms. Instead, LRMP and IRAG may interact with the distal N-terminus and/or distal C-terminus of HCN4, which are considerably longer than, and quite divergent from, the corresponding domains of other HCN channels. Isoform-specific regulation of HCN channels via non-conserved N- or C-terminal sequences is not without precedent; Filamin A interacts with HCN1 downstream of the CNBD at a site that is not conserved in other HCN isoforms leading to isoform-specific regulation of membrane trafficking (25).

Given the metabolic cost of producing large proteins, the long, unique termini of HCN4 are likely to be critical for isoform-specific channel function and regulation. In human HCN4, the non-conserved regions of the N- and C-termini distal to the HCN and CNBD together comprise 659 amino acids, more than 50% of the total sequence, and nearly double the length of corresponding regions in HCN1 or HCN2. Interestingly, an alternate transcriptional initiation site that removes the first 25 residues from HCN4 renders the channels insensitive to cAMP, despite the fact that cAMP binds to the C-terminus (51). Conversely, truncation of most of the HCN4 channel C-terminus at residue 719 restores cAMP sensitivity to HCN4 in CHO cells (14). And truncation of the non-conserved regions of the N- and C-termini of HCN4 shifts the voltage-dependence of channel activation to more hyperpolarized potentials in the absence of cAMP (52). Given this clear functional importance of the non-conserved regions of the HCN4 N- and C-termini, these domains represent rationale sites against which future pharmacological agents could be designed to specifically modulate HCN4 function, for example, to control heart rate (53).

### Physiological significance

HCN channels are key regulators of excitability in cells throughout the body. As one example of the physiological consequences of LRMP and IRAG modulation of HCN4 channels, our results support a potential role for IRAG in regulation of heart rate. It is well established that modulation of HCN4 and I_f_ in SAN myocytes — through phosphorylation, cyclic-nucleotides, pharmaceuticals, and mutations — alters heart rate and can cause bradycardia, tachycardia, and SAN dysfunction (7, 10–12, 15, 54). Our modeling predicts that IRAG expression and the ensuing depolarized V_1/2_ of I_f_ in the absence of cAMP increase the spontaneous AP firing rate in pacemaker cells, and thus contribute to a higher intrinsic heart rate. Indeed, this effect is in agreement with the limited data available from an IRAG knockdown mouse line, which has a lower resting heart rate (41). In the case of LRMP, our data suggest that expression in the SAN would limit increases in I_f_ in response to βAR stimulation. However, the only evidence at present for a role of LRMP in heart rate regulation is a GWAS study linking loci near LRMP to resting heart rate variability (37). Ultimately, the physiological roles of LRMP and IRAG in heart rate regulation will need to be determined by in-depth studies of sinoatrial myocytes, including those from the IRAG knockdown mouse line.

HCN4 and IRAG are also co-expressed at high levels in thalamocortical neurons (2, 55), where HCN4 is known to be an important regulator of input resistance, action potential burst firing, and thalamic and cortical oscillations during active wakefulness (2, 3). Although the phenotype of a brain-specific knockout of HCN4 is relatively mild, it is associated with a slowing of thalamic and cortical oscillations and possibly an increase in anxiety (2, 56). HCN4 loss-of-function is also implicated in generalized epilepsies that are characterized by electrical discharges that are believed to originate in thalamo-cortical circuits (57, 58). It will be interesting in future work to determine the role of IRAG modulation of HCN4 in thalamocortical neuron function.

### Macromolecular Complexes and ER-PM Junctions

As ER membrane proteins, LRMP and IRAG fall into an important and growing class of ER proteins that interact with ion channels on the plasma membrane. These include the ER transmembrane protein STIM1 that interacts with ORAI1 ion channels to mediate store-operated calcium release (59) and the junctophilins whose interactions with L-type calcium channels are critical for excitation-contraction coupling (60, 61). In SAN myocytes, the association between IRAG and HCN4 has significant implications as a potential physical link between two systems known to be important for spontaneous pacemaker activity: the plasma membrane HCN4 channels and the proteins that regulate calcium release from intracellular stores in the sarcoplasmic reticulum (62). Both LRMP and IRAG have previously been shown to associate with IP_3_Rs, which are ER Ca^2+^ release channels (42, 43). In SAN myocytes, calcium release through IP3 receptors has a downstream effect in regulating calcium release through ryanodine receptors (63), which in turn helps drive the diastolic depolarization of sinoatrial myocytes through the electrogenic sodium-calcium exchanger.

These findings also raise the possibility of large, macromolecular protein complexes involving HCN4 channels in SAN myocytes that may regulate pacemaker activity in response to nitric oxide (NO). Since IRAG in smooth muscle complexes with IP3 receptors and cGMP kinase Iβ to regulate the release of ER calcium through IP3 receptors in response to NO (42, 55), our data showing both functional and physical interactions between IRAG and HCN4 may portend a link between HCN4 channel activity and NO/cGMP/PKG signaling. The effects of NO on pacemaker function are not fully understood and are confounded by biphasic effects on pacemaker cells and effects on both branches of the autonomic nervous system (64–66). At the level of the SAN, NO leads to an increase in I_f_ and heart rate that is in part due to NO-stimulated increases in cGMP (66, 67). While, cGMP can directly bind the CNBD of HCN channels, it does so at approximately 10 fold lower affinity compared to cAMP (68). This suggests an interesting direction for future work examining a potential interaction of cGMP/PKG signaling with the proteins underlying sinoatrial pacemaking.

## Acknowledgments

The authors gratefully acknowledge Drs. Martin Biel and Eric Accili for providing reagents and Dr. Kika Sucharov for the use of her ABI 7300 Real-Time PCR System. The authors would also like to thank Dr. Christian Rickert for his input on data collection and analysis. Dr. Peters is funded by an American Heart Association Postdoctoral Fellowship (#19POST34380777). This work was supported by NIH R01 HL 088427 (to C.P.) and a grant from the Department of Physiology & Biophysics at the University of Colorado School of Medicine (to L.A.W. and C.P.).

## Materials and Methods

### Ethical approval and animals

This study was carried out in accordance with the US Animal Welfare Act and the National Research Council’s *Guide for the Care and Use of Laboratory Animals* and was conducted according to a protocol that was approved by the University of Colorado-Anschutz Medical Campus Institutional Animal Care and Use Committee (protocol number 84814(06)1E). Six-to eight-week old male C57BL/6J mice were obtained from Jackson Laboratories (Bar Harbor, ME; Cat. #000664). Animals were anesthetized by isofluorane inhalation and euthanized under anesthesia by cervical dislocation.

### Cell lines and DNA Constructs

Patch-clamp experiments were performed in HEK-293 cells (ATCC, Manassas, VA), an HCN4 stable line in HEK-293 (generously provided by Dr. Martin Biel’s lab) (69), or an HCN4 stable line in CHO-K1 cells (15). All cell lines were grown in a humidified incubator at 37°C and 5% CO2. HEK-293 cell lines were grown in high glucose DMEM with L-glutamine, supplemented with 10% FBS, 100 U/mL penicillin, and 100 μg/ml streptomycin (all from Thermo Fisher Scientific, Waltham, MA). To maintain the HEK-HCN4 stable line, the media was further supplemented with 200 μg/mL Hygromycin B (Invivogen, San Diego, CA). The CHO-K1 cells stably expressing HCN4 were grown in Ham’s F12 medium with L-glutamine (Thermo Fisher Scientific) supplemented with 10% FBS, 100 U/mL penicillin, 100 μg/ml streptomycin, and 100 μg/mL Zeocin (Invivogen). In CHO-K1 cells, HCN4 expression was induced by addition of 20 μg/mL of tetracycline to the growth media 24-48 h prior to experiments.

The mouse clone of LRMP in pCMV6 was purchased from OriGene (Rockville, MD; Cat. #MC228229). The mouse variant of IRAG in pReceiver-M61 was purchased from GeneCopoeia (Rockville, MD; Cat. #EX-Mm30453-M61). All HCN4 experiments were performed in stable cell lines. Patch-clamp experiments on HCN1 and HCN2 were performed with transient transfection of 2 μg of either pCDNA3.1-HCN1 or pCDNA3.1-HCN2 (generously provided by Dr. Eric Accili) in HEK293 cells using Fugene6 (Roche, Basal, Switzerland) according to the manufacturer’s instructions. Two μg of either LRMP or IRAG was cotransfected for patch-clamp experiments. All transfections, except for those including IRAG, which has an IRES-eGFP marker, were performed with the addition of 0.5 μg of eGFP as a marker. For western-blotting and co-immunoprecipitation experiments, 15 μg of LRMP or IRAG was transfected using electroporation with a Lonza 4D-Nucleofecter (Basal, Switzerland) according to the manufacturer’s instructions.

For co-immunoprecipitation experiments a Myc-tag was added to the N-terminal end of LRMP or IRAG immediately following the initial methionine. A C-terminal Myc-tagged IRAG clone did not functionally regulate HCN4 (data not shown), and the C-terminal Myc-tag in LRMP appeared to be cleaved off (data not shown). The latter is likely due to a previously described post-translational modification that removes the luminal domain of LRMP (39). As the transfection efficiency of the pReciever-M61 IRAG vector was low, the Myc-IRAG clone was sub-cloned into pCDNA3.1 for co-immunoprecipitation and western blotting experiments.

### LRMP and IRAG knockdown in CHO cells

To knockdown endogenous LRMP or IRAG expression in CHO cells we used a CRISPR Cas9 system (70). LRMP- and IRAG-specific gRNA sequences were cloned into the pSpCas9(BB)-2A-GFP vector (Addgene, Watertown, MA; Cat. #PX458). Two distinct cut sites were used for each IRAG (XM_027410089.1) and LRMP (XM_027430141.1). Primer sequences are presented in the SI materials.

LRMP and IRAG knockdowns were performed by transient transfection of both of the appropriate CRISPR plasmids into a CHO cell line stably expressing HCN4. To exclude an effect of the CRISPR protein itself, control experiments were performed on CHO cells stably expressing HCN4 transfected with the pSpCas9(BB)-2A-GFP vector lacking a gRNA sequence.

### Patch-Clamp Electrophysiology

For patch-clamp experiments, cells were plated on sterile, protamine-coated glass coverslips 24-48 hours prior to experiments. Cells were transferred to a glass-bottom recording chamber and perfused (~1 ml/min) at room temperature with extracellular recording solution containing (in mM): 30 KCl, 115 NaCl, 1 MgCl_2_, 1.8 CaCl_2_, 5.5 glucose, and 5 HEPES. Transfected cells were identified by green fluorescence.

Patch clamp recordings used pulled borosilicate glass pipettes with resistances of 1.0-2.5 MOhm when filled with intracellular solution containing (in mM): 130 K-aspartate, 10 NaCl, 1 EGTA, 0.5 MgCl_2_, 5 HEPES, and 2 MgATP. 1 mM cAMP was added to the intracellular solution for some experiments as indicated. Data were acquired at 5 KHz and low pass filtered at 10 KHz using an Axopatch 200B amplifier, Digidata 1440A A/D converter, and Clampex software (Molecular Devices, San Jose, CA). The fast capacitance component, corresponding to pipette capacitance was compensated in all experiments. Membrane capacitance and series resistance (Rs) were estimated in whole-cell experiments using 5 mV test pulses. Only cells with Rs < 10 MOhm were analyzed. All data were analyzed in Clampfit 10.7 (Molecular Devices).

Channel activation was estimated from the peak tail current at −50 mV following 3 s hyperpolarizations to membrane potentials between −50 mV and −170 mV from a holding potential of 0 mV. Peak tail currents were fit by a single Boltzmann curve to yield midpoint activation voltages (V½) and slope factors. Channel activation rates were assessed by measuring the time to half peak current during hyperpolarizing pulses to −150 mV. Channel deactivation rates were assessed by fitting exponential curves to the decay of tail currents at −50 mV following hyperpolarizing pulses to between −130 mV and −150 mV. All experiments and values are corrected for a calculated +14 mV junction potential.

### qPCR

Isolated SAN or left atrial tissue was homogenized using a bead homogenizer and total RNA was extracted using QIAzol (Qiagen, Hilden, Germany) and chloroform according to the manufacturer’s instructions. RNA samples from heart tissue and HEK and CHO cell pellets were purified using a RNeasy mini kit (Qiagen). RNA was reverse-transcribed to cDNA using an Applied Biosystems High-Capacity RNA-to-cDNA kit (Foster City, CA) according to the manufacturer’s instructions. Two independent reverse transcription reactions were performed for each RNA sample. qPCR experiments were performed on an ABI 7300 Real-Time PCR System (Applied Biosystems) using Applied Biological Materials’ BrightGreen qPCR master mix kit or New England BioLabs’s Luna Universal qPCR master mix (Ipswich, MA) according to the manufacturer’s instructions. Primer sequences are listed in the SI materials.

qPCR primers were designed to span exon-exon junctions to prevent replication of contaminating genomic DNA. qPCR reactions for each cDNA prep were run at 5 different cDNA concentrations between 0.001 ng and 10.0 ng with each primer set. Efficient cDNA doubling was confirmed by comparing Cq values from control 18S ribosomal RNA reactions across concentrations. No-template controls were run for each cDNA-primer combination to confirm that contamination was not present. Cq values for LRMP, IRAG, and HCN4 were normalized to 18S Cq values from the same cDNA prep at each template concentration.

### Western Blotting

Isolated SAN, left atrial, ventricular, and liver tissues were homogenized using a dounce homogenizer in a modified radioimmunoprecipitation assay (RIPA) buffer (150 mM NaCl, 25 mM Tris, 1% NP40, 1% sodium deoxycholate, and 1% SDS) with Roche protease inhibitors. Cells were lysed at 4°C for 3 h after which genomic DNA and cellular fragments were pelleted. Protein concentration was assessed using A280 absorbance on a NanoDrop spectrophotometer (Thermo Fisher Scientific). Samples were mixed with 2X Laemelli’s sample buffer and heated to 60°C for 30 min before loading. HEK-293 and CHO-K1 cell samples were prepared in the same manner without the dounce homogenization step.

Electrophoresis was performed using a BioRad Mini-PROTEAN cell with 4-12% or 4-15% precast TGX gels (BioRad, Hercules, CA) in TGS running buffer. Transfer to membranes was performed in BioRad Mini Trans-Blot cells in TGS buffer with 20% methanol or in a BioRad Trans-Blot Turbo in Bjerrum Schafer-Nielsen buffer. Following transfer, membranes were blocked for 1 h in 5% W/V nonfat dry milk at room temperature. Primary antibodies were applied overnight at 4°C in 5% milk. Following washing, secondary antibodies were applied at room temperature for 1 h in 5% milk. Chemiluminescence was measured using Immobilon Western HRP Substrate (Millipore, Burlington, MA) on a Kodak Digital Science Image Station 440CF (Kodak, Rochester, NY) or on a Li-Cor Odyssey Fc imaging system (LI-COR Biosciences, Lincoln, NE). Protein concentration in all samples was quantified using A280 absorbance prior to loading, and GAPDH was used as a loading control.

For co-immunoprecipitation experiments cells were lysed at 4°C for 30 min in a buffer containing 120 mM NaCl, 50 mM Tris (pH 7.4), 1 mM EDTA, 0.25% deoxycholate and 1% NP-40 with Roche protease inhibitors. Following pelleting of genomic DNA and cellular fragments, lysates (250 μg) were incubated overnight at 4°C with a 1:100 dilution of anti-HCN4 antibody (Alomone Labs, Jerusalem; Cat. #APC-052). Antibody-HCN4 complexes were immunoprecipitated with protein A/G PLUS-agarose (Santa Cruz; Cat. #SC-2003) and washed 1x with lysis buffer and 2x with PBS. Protein was eluted in 1X Laemelli’s sample buffer at 60°C for 10min. Proteins were separated on 4-15% gradient gels (BioRad). For whole extract and the supernatant fraction after immunoprecipitation, 10 μg protein was loaded. For the immunoprecipate, half of the eluate volume was loaded. Proteins were electrophoresed for 1.5 hours at 150 V and transferred to PVDF membranes at 100 V for 1 hour. Membranes were blocked with 5% BSA for 1 hour and western blotted with anti-Myc antibody (Invitrogen; MA1-21316) overnight at 4 °C. Membranes were washed 3X with tris-buffered saline with 0.05% Tween-20 (TBST) and incubated with anti-mouse secondary antibody (Sigma) at 1:50,000 for 1 hour at RT. After washing, membranes were visualized using enhanced chemiluminescence and film exposures of varying lengths.

Antibodies used for western blotting were: mouse anti-Myc.A7 (Thermo Fisher Scientific; Cat. #MA1-21316) at 1:1,000 to 1:5,000 dilution; mouse anti-HCN4 (NeuroMab, Davis, CA; Cat. #N114/10) at a 1:1,000 dilution; rabbit anti-HCN4 (Alomone Labs; Cat. #APC-052) at a 1:1,000 dilution; rabbit anti-IRAG (Thermo Fisher Scientific; Cat. #PA5-32111) at 1:500 to 1:1,000 dilution; mouse anti-GAPDH 6C5 (Thermo Fisher Scientific; Cat. #AM4300) at 1:2,500 to 1:5,000 dilution; goat anti-rabbit IgG HRP (Thermo Fisher Scientific; Cat. #31460) at a 1:1,000 dilution; and goat anti-mouse IgG HRP (Thermo Fisher Scientific; Cat. #31430) at a 1:10,000 dilution. We also tested two commercially-available LRMP antibodies: a rabbit anti-LRMP C-terminal antibody (Abgent, San Diego, CA; Cat. #AP19787b) and a rabbit anti-LRMP aa354-403 antibody (LSBio, Seattle, WA; Cat. #LS-C145883), but found that neither showed specific staining of LRMP. We also evaluated rabbit anti-IRAG (Thermo Fisher Scientific; Cat. #PA3-851) but found it to be less specific than PA5-32111.

### Statistics

All statistical analysis was performed using JMP Pro 14 software (SAS Institute, Cary, NC). Comparisons of patch-clamp data from LRMP- or IRAG-transfected, HCN1, 2, or 4-expressing cells were compared to untransfected cells using a two-factor ANOVA with the presence/absence of LRMP or IRAG and the presence/absence of cAMP as main factors. A differential effect of cAMP in the presence/absence of LRMP or IRAG was analyzed in the statistical model as an interaction between the two factors. A two-factor ANOVA with the same main factors and interaction factor was used to compare CHO cells transfected with CRISPR alone to those transfected with CRISPR and gRNAs for LRMP or IRAG. For qPCR data, a one-factor ANOVA with the measured gene as the main factor was used to test for a difference in sinoatrial LRMP and IRAG expression compared to HCN4 transcript expression. Student’s T-tests were used to compare IRAG or LRMP expression between HEK and CHO cells. All time constant values were log-transformed prior to statistical analysis because log-transformed time constants are normally distributed. Statistical significance was evaluated at P < 0.05.

All means, standard errors, and N values for patch-clamp recordings are provided in Supplementary Tables S2-4.

### I_f_ Models

Our I_f_ model was based on that proposed by Verkerk and Wilders which uses a single activation gate (Fig. 6C) that controls channel opening and closing (46). Equations used to model the activation gate are:

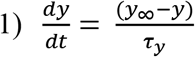

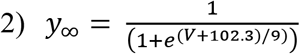

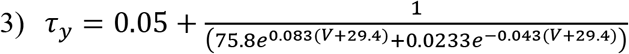

Where y is the activation gate, y_∞_ is the voltage-dependent steady-state value of the activation gate, τ_y_ is the voltage-dependent time constant of the activation gate, and V is membrane potential. The model of I_f_ uses a conductance of 0.224 pS/pF and a reversal potential of −34.8 mV. The wild-type current model uses the curves proposed by Verkerk and Wilders (46) that have been shifted along the voltage axis by −29.4 mV to match our previously recorded data (47). To delineate a possible range of I_f_ values, the IRAG over-expression model shifts both the time constant and steady-state curves to 7.6 mV more depolarized, the most depolarized we have observed the I_f_ V_1/2_ value with maximal isoproterenol stimulation (47). The IRAG knockout model shifts both the time constant and steady-state curves by 6 mV in the hyperpolarizing direction. These values were chosen to equal a total range of 13.6 mV between the IRAG knockout model and IRAG over-expression model (47), which is the range of values we observed for HCN4 in HEK cells between basal and cAMP stimulated conditions (Fig. 2). I_f_ currents from all three models were stimulated using either a 3 s square hyperpolarizing pulse to recapitulate the activation voltage dependence or a mouse sinoatrial myocyte action potential voltage waveform recorded within our lab. All calculations were performed in Python 3.7 using a forward Euler method with a 200 μs time step.

