## Supplemental Information for "Isoform-specific regulation of HCN4 channels by a family of endoplasmic reticulum proteins"

**Supporting Information**

*SI Methods*

CRISPR Primers

Note that the initial “G” is not part of the gene sequence and is used to facilitate expression

IRAG KO 1

G-148-CTGAGGATGAGGAGCCCCC-166

IRAG KO2

G-2276-GAGCCAGGAGACCAAGGCC-2294

LRMP KO1

G-584-AGCGAGGACCTTAGTGCAG-602

LRMP KO2

G-464-CTGTGTGAAGATGACAACC-482

qPCR Primers:

Human 18s Forward Primer: GGCCCTGTAATTGGAATGAGTC

Human 18s Reverse Primer: CCAAGATCCAACTACGAGCTT

Mouse 18s Forward Primer: GCAATTATTCCCCATGAACG

Mouse 18s Reverse Primer: GGCCTCACTAAACCATCCAA

Chinese hamster 18s Forward Primer: CTGAGAAACGGCTACCACATC

Chinese hamster 18s Reverse Primer: GCCTCGAAAGAGTCCTGTATTG

Human HCN4 Forward Primer: CTGAGAACTGGAAGGACTTAGC

Human HCN4 Reverse Primer: CAGGACAAGACTGTGGGTTT

Mouse HCN4 Forward Primer: GGTGGAGGACAACACAGAAA

Mouse HCN4 Reverse Primer: CAGGGATGGAGGAGATGAAATC

Human LRMP Forward Primer: AGTGGGATGTCTCTTCAGTTTATG

Human LRMP Reverse Primer: AGAGCCAGAGGGCCTTATTA

Mouse LRMP Forward Primer: CTGTACCAGAATGAACGAGGAC

Mouse LRMP Reverse Primer: GACGATGGCTGTGTGAGATAC

Chinese Hamster LRMP Forward Primer: TTCCAAGCCATCGTCTCTTC

Chinese Hamster LRMP Reverse Primer: CCGGCTGAACCTGTCATTAT

Human IRAG Forward Primer: CTGAGGAACCTGAAGAGGTAGA

Human IRAG Reverse Primer: GCTGACACAGTTTGGGATACA

Mouse IRAG Forward Primer: GTTAACCCACCTCTTGGGTAAG

Mouse IRAG Reverse Primer: GGTCTGTGGAAGGACACATTAG

Chinese Hamster IRAG Forward Primer: CTCCCATAACTCACCCTGAATAG

Chinese Hamster IRAG Reverse Primer: CCTCACTCTGAACCTCATTACC

*Supplementary Table S1: Significantly greater endogenous LRMP and IRAG mRNA Transcript Levels in CHO cells compared to HEK cells.*

| Gene | CHO HCN4 Norm Cq | HEK HCN4 Norm Cq | Fold Difference CHO vs HEK |
|---|---|---|---|
| LRMP | -20.9 ± 0.7 (3) | -25.4 ± 0.3 (3) | 20.0 ± 2.3 |
| IRAG | -20.3 ± 0.3 (3) | -24.1 ± 0.1 (3) | 14.6 ± 1.4 |

Cq value is the PCR cycle during which the fluorescence passes the detection threshold. All Cq values were normalized to the Cq value for 18S primers from the same biological sample and technical replicate to control for any errors in the initial RNA quantitation. As the expression of 18S is greater than LRMP or IRAG, the normalized values are negative. A less negative Cq value represents fewer doubling steps to pass the detection threshold and therefore a greater initial quantity of cDNA. Values are reported as mean ± SEM. Number of independent biological samples are reported in parenthesis.

*Supplementary Table S2: Properties of HCN4 currents in HEK cells in the presence or absence of LRMP or IRAG.*

| | **Activation Midpoint (mV)** | | **ΔV½ (mV)** | **Slope factor** | | **Current Density (pA/pF)** | |
|---|---|---|---|---|---|---|---|
| | **No cAMP** | **1 mM cAMP** | | **No cAMP** | **1 mM cAMP** | **No cAMP** | **1 mM cAMP** |
| HCN4 | -116.3 ± 1.0 (22) | -103.2 ± 1.9 (16) | 13.1 | 10.9 ± 0.3 (22) | 13.1 ± 0.8 (16) | 138 ± 14 (22) | 124 ± 19 (15) |
| HCN4 + LRMP | -114.4 ± 1.2 (15) | -109.3 ± 1.6 (15) | 5.1 | 10.1 ± 0.4 (15) | 13.8 ± 0.5 (15) | 118 ± 18 (14) | 152 ± 23 (13) |
| HCN4 + Myc-LRMP | -113.1 ± 1.6 (10) | -108.0 ± 1.7 (11) | 5.1 | ___ | ___ | ___ | ___ |
| HCN4 + IRAG | -109.2 ± 1.8 (20) | -105.7 ± 1.9 (14) | 5.0 | 11.3 ± 0.3 (20) | 13.9 ± 0.5 (14) | 144 ± 27 (17) | 175 ± 35 (11) |
| HCN4 + Myc-IRAG | -105.0 ± 1.6 (11) | -101.4 ± 2.3 (6) | 3.9 | ___ | ___ | ___ | ___ |

Values are reported as mean ± SEM. Number of independent data points are reported in parenthesis. Activation midpoints from Boltzmann fits to conductance-voltage data. Current density was measured in response to 3 s steps to -150 mV.

*Supplementary Table S3: Time Constants of HCN4 Activation and Deactivation in the Presence or Absence of 1 mM cAMP in HEK cells.*

| | **Time to half activation @ -150mV (ms)** | | **$\tau_{Deactivation}$ -50mV (ms)** | |
|---|---|---|---|---|
| | **No cAMP** | **1 mM cAMP** | **No cAMP** | **1 mM cAMP** |
| HCN4 | 372 ± 17 (22) | 210 ± 15 (16) | 692 ± 33 (22) | 1002 ± 58 (15) |
| HCN4 + LRMP | 324 ± 24 (14) | 248 ± 16 (12) | 781 ± 49 (15) | 970 ± 70 (13) |
| HCN4 + IRAG | 294 ± 30 (15) | 218 ± 18 (11) | 763 ± 43 (20) | 1221 ± 115 (12) |

All values are reported as mean ± SEM. Number of independent data points are reported in parenthesis.

*Supplementary Table S4: HCN Channel midpoints in the presence or absence of LRMP or IRAG.*

| **Cell Type, Channel** | **Transfection** | **Activation Midpoint (mV)** | | **ΔV½ (mV)** |
|---|---|---|---|---|
| | | **No cAMP** | **1 mM cAMP** | |
| HEK - HCN1 | — | -75.2 ± 1.9 (10) | -74.1 ± 3.5 (6) | 1.1 |
| | LRMP | -77.5 ± 1.0 (8) | -76.5 ± 1.3 (6) | 1.0 |
| | IRAG | -78.1 ± 1.5 (8) | -74.8 ± 2.3 (5) | 3.3 |
| HEK - HCN2 | — | -103.8 ± 0.9 (7) | -80.8 ± 1.1 (5) | 23.0 |
| | LRMP | -102.0 ± 0.7 (8) | -81.2 ± 1.1 (9) | 20.8 |
| | IRAG | -103.9 ± 1.4 (6) | -82.7 ± 3.5 (5) | 21.2 |
| CHO - HCN4 | — | -109.7 ± 1.3 (12) | -108.1 ± 1.7 (14) | 1.6 |
| | CRISPR | -112.9 ± 1.9 (13) | -111.1 ± 1.8 (10) | 1.8 |
| | LRMP KO | -120.5 ± 1.9 (13) | -107.3± 2.3 (13) | 13.2 |
| | IRAG KO | -114.8 ± 1.8 (15) | -110.2 ± 1.2 (15) | 4.6 |

All values are reported as mean ± standard error of the mean. Number of independent data points are reported in parenthesis.

**Figure S1**

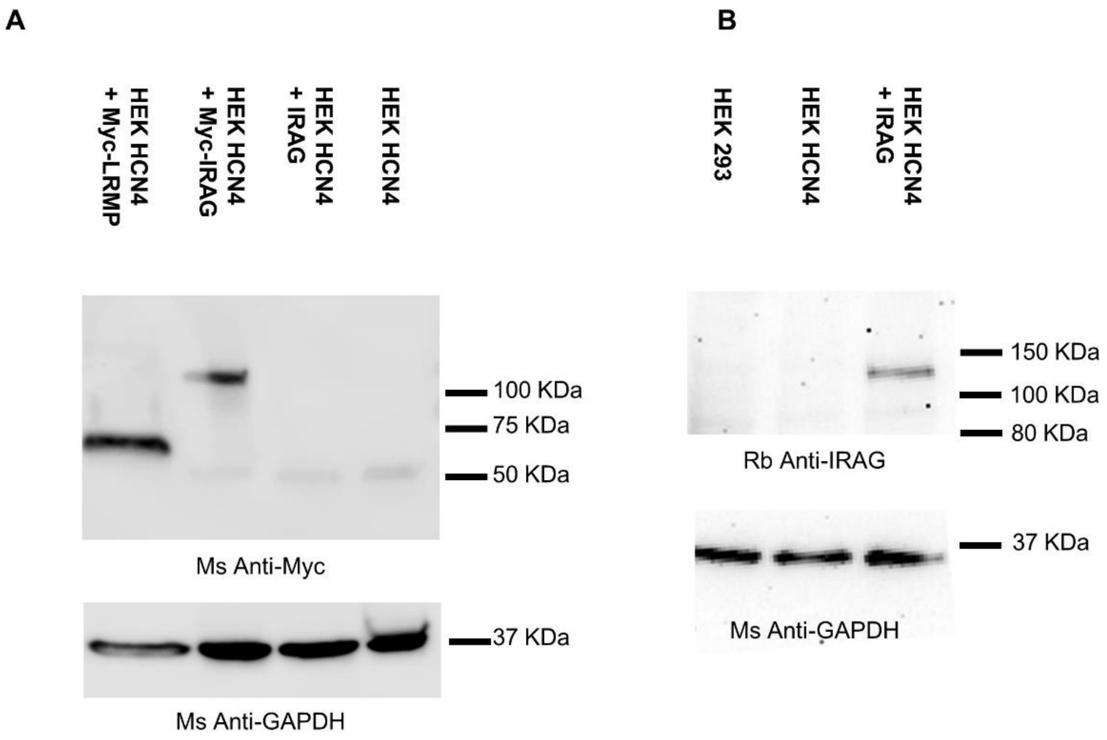


*Figure S1: Anti-Myc and anti-IRAG antibodies are specific for their target proteins.*

**A:** Western blot of anti-Myc antibody staining in HEK cells stably expressing HCN4 in the absence or presence of IRAG, Myc-tagged IRAG, or Myc-tagged LRMP. GAPDH staining is used as a loading control. The faint band seen at 50 kDa is likely endogenous C-Myc protein. **B:** Western blot of anti-IRAG antibody staining in HEK cell lysates in the absence or presence of HCN4 and IRAG. GAPDH staining is used as a loading control.

**Figure S2**

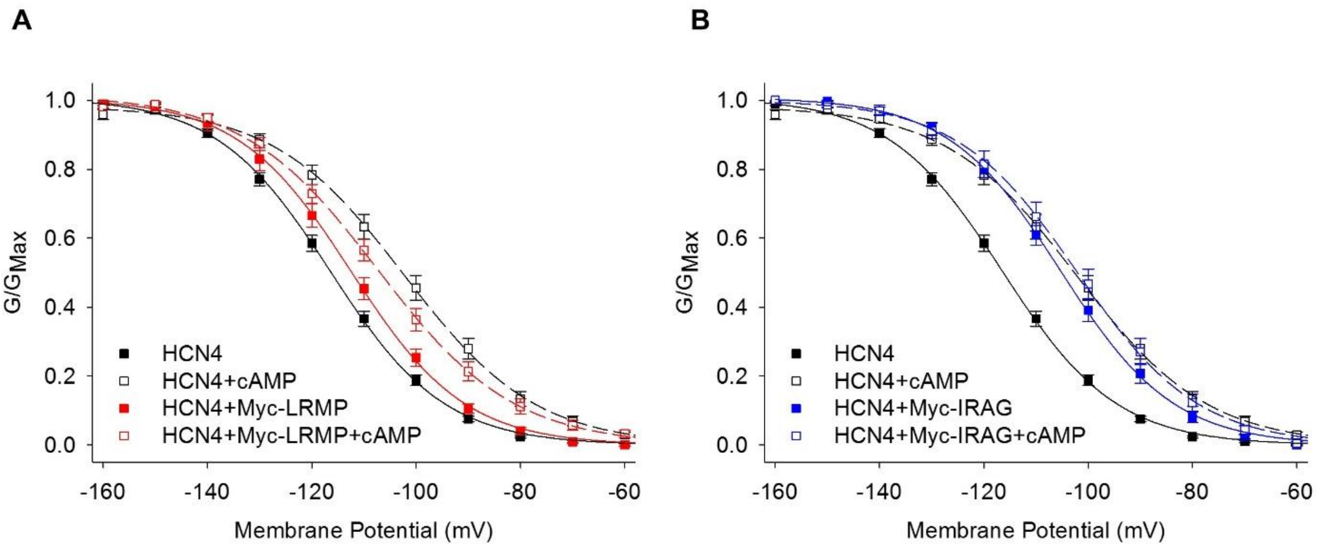


*Figure S2: Addition of an N-terminal Myc-tag does not impair the ability of LRMP or IRAG to regulate HCN4 gating.*

**A, B:** Average conductance-voltage relations for HCN4 in control conditions (*black*), the presence of Myc-LRMP (*red*), or the presence of Myc-IRAG (*blue*). GVs in the presence of 1 mM cAMP are shown by *open symbols*. Error bars are SEM, N = 6-11 (See Table S2). Control HCN4 data in both panels are the same as those in Fig. 2.
